# Mitochondrial calcium uptake regulates tumour progression in embryonal rhabdomyosarcoma

**DOI:** 10.1101/2021.11.10.468020

**Authors:** Hsin Yao Chiu, Amos Hong Pheng Loh, Reshma Taneja

## Abstract

Embryonal rhabdomyosarcoma (ERMS) is characterized by a failure of cells to complete skeletal muscle differentiation. Although ERMS cells are vulnerable to oxidative stress, the relevance of mitochondrial calcium homeostasis in oncogenesis is unclear. Here, we show that ERMS cell lines as well as primary tumours exhibit elevated expression of the Mitochondrial Calcium Uniporter (MCU). MCU knockdown resulted in impaired mitochondrial calcium uptake and a reduction in mitochondrial reactive oxygen species (mROS) levels. Phenotypically, MCU knockdown cells exhibited reduced cellular proliferation and motility, with an increased propensity to differentiate *in vitro* and *in vivo*. RNA-sequencing of MCU knockdown cells revealed a significant reduction in genes involved in TGFβ signalling that play prominent roles in oncogenesis and inhibition of myogenic differentiation. Interestingly, modulation of mROS production impacted TGFβ signalling. Our study elucidates mechanisms by which mitochondrial calcium dysregulation promotes tumour progression and suggests that targeting the MCU complex to restore mitochondrial calcium homeostasis could be a therapeutic avenue in ERMS.

## Introduction

Rhabdomyosarcoma (RMS) is the most prevalent soft-tissue sarcoma in childhood and adolescence [1–3]. Even though RMS cells express myoblast determination protein 1 (MYOD), a master regulator of myogenic differentiation, they exhibit a failure to complete the differentiation program. The two main subtypes are embryonal rhabdomyosarcoma (ERMS) and alveolar rhabdomyosarcoma (ARMS) that account for approximately 70% and 20% respectively of all RMS cases [1–4]. ERMS cells possess a more complex karyotype with a loss of heterozygosity at 11p15.5 and a higher mutation burden compared to ARMS [4,5]. Mutations in RAS, receptor tyrosine kinase or phosphoinositide-3 kinase (PI3K) complexes are most commonly found in ERMS [2,5]. These pathways maintain redox balance and energy metabolism for cellular functions [6,7]. Given the genetic aberrations in ERMS and the importance of mitochondrial function in cancer, a few studies have demonstrated that reactive oxygen species (ROS) production and cellular metabolism are altered in ERMS [8–10]. Upregulation of mitochondrial genes in patient tumours has also been reported [11]. These observations suggest that mitochondrial dysfunction may be important in ERMS oncogenesis. Nevertheless, the role of mitochondrial calcium (Ca^2+^) homeostasis has not been characterised.

Mitochondrial calcium uniporter (MCU) complex is the main channel responsible for mitochondrial Ca^2+^ uptake and requires inner mitochondrial membrane (IMM) potential for Ca^2+^ to enter the mitochondrial matrix. The MCU complex plays a fundamental role in regulating global Ca^2+^ signalling, redox balance, aerobic metabolism and apoptosis [12,13]. Mitochondrial calcium uniporter (MCU) is the main pore-forming protein. The loss of MCU inhibits mitochondrial Ca^2+^ uptake by approximately 75% [14,15]. Mitochondrial calcium uptake 1 (MICU1) is the gatekeeper of MCU and forms a heterodimer with MICU2 [16,17]. The MICU1-MICU2 complex prevents mitochondrial Ca^2+^ overload under basal cytosolic Ca^2+^ conditions. MICU1 acts to regulate the threshold of MCU opening and cooperates with MICU2 to activate the channel under high Ca^2+^ concentration [18].

MCU and MICU1 deregulations have been reported in several cancers [13,19,20]. For instance, MCU overexpression in breast cancer correlates with tumour size, invasiveness and poor prognosis [21]. In colorectal cancer, MCU-induced mitochondrial Ca^2+^ uptake promotes mitochondrial biogenesis and tumour growth through mitochondrial transcription factor A (TFAM) and Nuclear factor kappa-light-chain-enhancer of activated B cells (NF-κB) [22]. The elevation in mitochondrial Ca^2+^ through MCU overexpression promotes hepatocellular carcinoma (HCC) metastasis through ROS production [23]. In contrast, the downregulation of MCU in cervical and colon cancer favours survival [24]. MICU1 expression is deregulated in liver, breast and ovarian cancer. Low MICU1 expression in HCC is correlated with poor prognosis [25], but paradoxically MICU1 overexpression in ovarian cancer correlates with poor survival and chemoresistance [26,27]. While the deregulation of their expressions varies in cancer-type specific manner, in general, overexpression of MCU and loss of MICU1 expression is correlated with poor prognosis [13,28].

In this study, we show that MCU is overexpressed in ERMS tumours and its silencing causes a reduction in mitochondrial Ca^2+^ uptake. This is correlated with reduced mitochondrial ROS (mROS) production. MCU reduction impaired cellular proliferation and motility, while enhancing myogenic differentiation. Interestingly, transforming growth factor beta (TGFβ) signalling pathway was dampened upon MCU knockdown consequent to reduced mROS levels. Elevating mROS reversed the phenotypes observed upon MCU depletion. Our study elucidates the relevance of mitochondrial Ca^2+^ signaling in driving tumour progression.

## Results

### MCU is overexpressed in ERMS

Previous studies have suggested deregulation of oxidative stress in ERMS [10,29,30]. We therefore examined mitochondrial Ca^2+^ uptake, oxygen consumption rate (OCR) and adenosine triphosphate (ATP) production in three patient derived ERMS cell lines (RD, RD18 and JR1). As controls, we used primary human skeletal muscle myoblasts (HSMM) and two ARMS cell lines (RH30 and RH41). Live cell staining using Rhod2-AM revealed significantly elevated mitochondrial Ca^2+^ in ERMS cell lines as compared to HSMM and ARMS cell lines (Fig. 1A, left panel). The uptake of mitochondrial Ca^2+^ upon induction is crucial in relaying signals. Basal fluorescence was measured for 1min before Ca^2+^ uptake was induced with 100μM of histamine. Maximal mitochondrial Ca^2+^ uptake was measured by quantifying the difference between basal fluorescence and highest fluorescence intensity attained post-induction. A significant increase in basal and maximal mitochondrial Ca^2+^ uptake was seen in all three ERMS cell lines relative to HSMM (Fig. 1A, right panels). We then analysed mitochondrial function through measurement of oxygen consumption rate (OCR) and ATP production. A significant increase in basal and maximal OCR as well as in ATP production were seen in ERMS cell lines relative to HSMM and ARMS cell lines (Fig. 1B). Together, these results demonstrate that ERMS cell lines exhibit dysregulated mitochondrial functions.

**Fig. 1.**
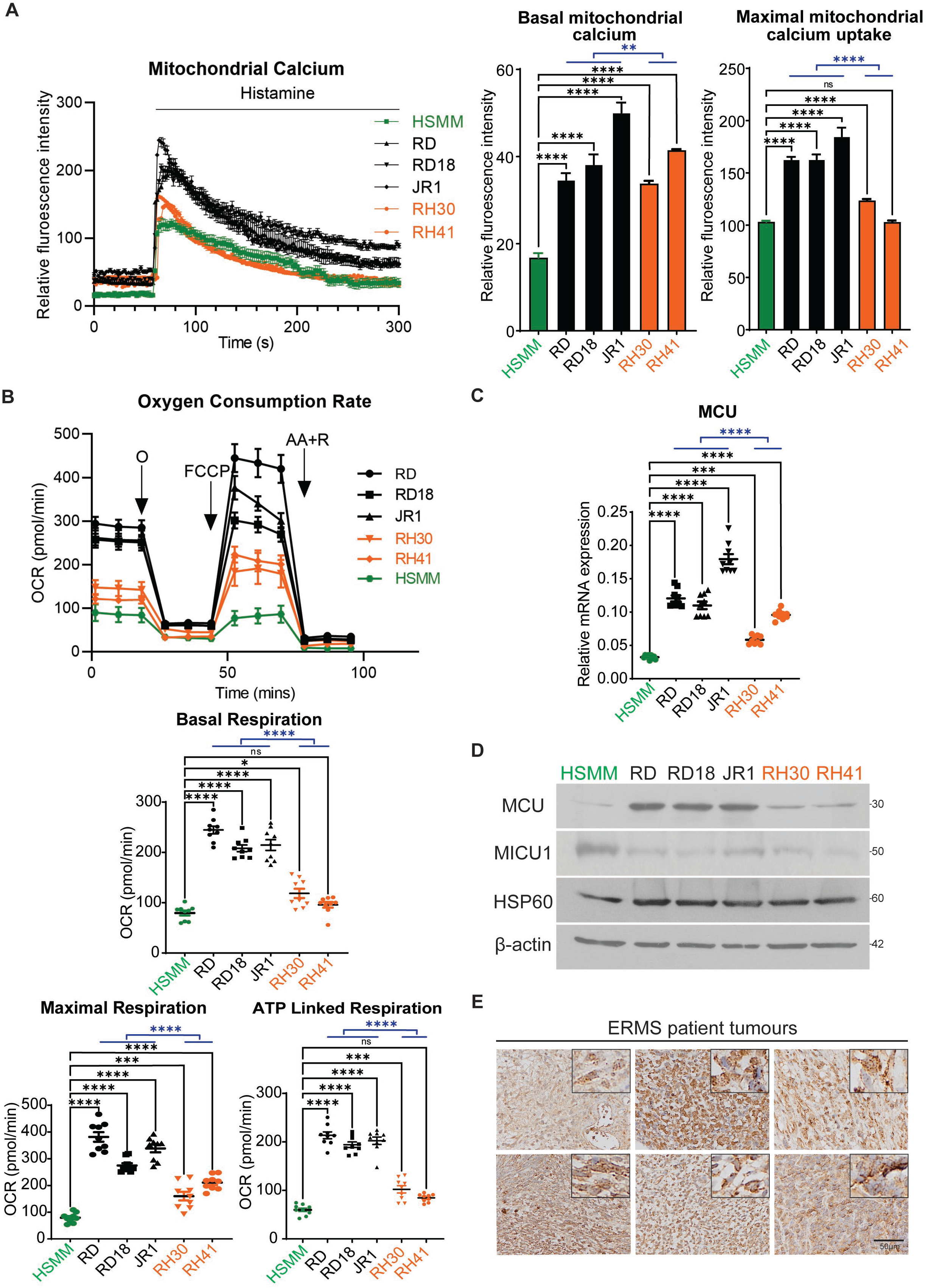
Altered mitochondrial function and MCU expression in ERMS cell lines and patient samples. A. Basal and maximal mitochondrial Ca^2+^ levels in HSMM, RD, RD18, JR1, RH30 and RH41 cells was measured with Rhod2-AM staining. The graph on the right below shows basal and maximal mitochondrial Ca^2+^ uptake upon induction with 100μM histamine (n=3). Values correspond to the average ± SEM. The blue line indicates significance calculated by comparing the average of ERMS cell lines and ARMS cell lines. (ns=not significant, ** *p*≤0.01 and **** *p*≤0.0001). B. Oxygen consumption rate (OCR) was measured with Seahorse analyser with the addition of O: Oligomycin, FCCP: Carbonyl cyanide-4-(trifluoromethoxy)-phenylhydrazone, AA+R: Antimycin A and Rotenone were added accordingly. Values correspond to average ± SEM (n=3). Basal and maximal respiration rate of HSMM, ERMS and ARMS cell lines as well as mitochondrial ATP production are shown (n=3). The blue line shows significance between the average of ERMS cell lines and ARMS cell lines. (* *p*≤0.05, *** *p*≤0.001 and **** *p*≤0.0001). C. MCU mRNA was examined in HSMM, RD, RD18, JR1, RH30 and RH41 by qPCR analysis (n=3). Values correspond to average ± SEM. Statistical significance was calculated by one-way ANOVA analysis. The blue line shows the significance comparing the average of ERMS cell lines and ARMS cell lines. (** *p*≤0.01, *** *p*≤0.001, **** *p*≤0.0001). D. Western blot analysis showing MCU, MICU1 and HSP60 protein levels in HSMM, RD, RD18, JR1, RH30 and RH41 cells. β-actin was used as loading control. A representative image of 3 independent experiments is shown. E. 6 archival ERMS patient tumour specimens were analysed by IHC using anti-MCU antibody. Images were taken at 40X magnification. Inset shows 3X zoomed in image. Scale bar: 50μm.

Mitochondrial Ca^2+^ uptake into the inner mitochondrial matrix is tightly regulated by the MCU complex. Therefore, we examined the expression of MCU, the main pore forming subunit of MCU complex in ERMS. MCU was found to be overexpressed in all three ERMS cell lines at both the mRNA and protein level compared to HSMM and ARMS cell lines (Fig. 1C and Fig. 1D). On the other hand, MICU1 expression was downregulated in all three ERMS cell lines (Fig. 1D). No significant difference was observed in the expression Heat Shock Protein 60 (HSP60) a mitochondrial molecular chaperone (Fig. 1D), suggesting that there was no overt change in mitochondrial mass that would account for the change in MCU and MICU1 expression. MCU expression was also examined in six archival ERMS tumour sections by immunohistochemistry (IHC) using anti-MCU antibody (Fig. 1E). All samples showed high MCU expression with varying degrees of speckling. Similarly, tissue microarray (TMA) of 27 ERMS patient tumours showed elevated MCU expression as compared to 24 ARMS patient samples and 8 normal muscles (Supplementary Fig. 1).

### MCU upregulation regulates mitochondrial function

To investigate the relevance of MCU overexpression, RD cells were transfected with empty vector (shScr) or MCU-specific shRNA (shMCU). The knockdown of MCU was specific with no change in MICU1, MICU2 and HSP60 levels (Fig. 2A). We next examined the effect of MCU knockdown on mitochondrial Ca^2+^ concentration using Rhod2-AM which localised specifically in the mitochondria as seen by co-localisation with MitoTracker (Fig. 2B). Upon histamine induction, a pronounced 65% reduction in maximal mitochondrial Ca^2+^ uptake was observed in shMCU cells with a small, albeit significant decrease in basal mitochondrial Ca^2+^ (Fig. 2B). A significant reduction in MitoSOX staining as well as reduced cytosolic ROS was seen with CM-H_2_DCFDA staining in shMCU cells (Fig. 2C-D). Since ATP and mROS are produced by electron transport chain (ETC) during cellular respiration, we examined ATP production and OCR. A significant reduction in overall ATP production was observed in shMCU cells (Fig. 2E). Correspondingly, up to 70% reduction in basal and maximal respiration rate was seen upon MCU knockdown and ATP-linked respiration through oxidative phosphorylation (OXPHOS) also showed a significant reduction (Fig. 2F). Consistent with stable MCU knockdown in RD cells, transient MCU knockdown in JR1 cells showed significant reduction in maximal mitochondrial Ca^2+^ uptake with no change in basal mitochondrial Ca^2+^ (Supplementary Fig. 2A-B). In addition, a significant reduction in MitoSOX staining was observed in siMCU cells (Supplemntary Fig. 2C). Similarly, a significant decrease in basal and maximal respiration rate as well as ATP-linked repiration upon MCU knockdown (Supplementary Fig. 2D-E). Together these results demonstrate that modulation of MCU expression is sufficient to change mitochondrial function in ERMS cell lines.

**Fig. 2.**
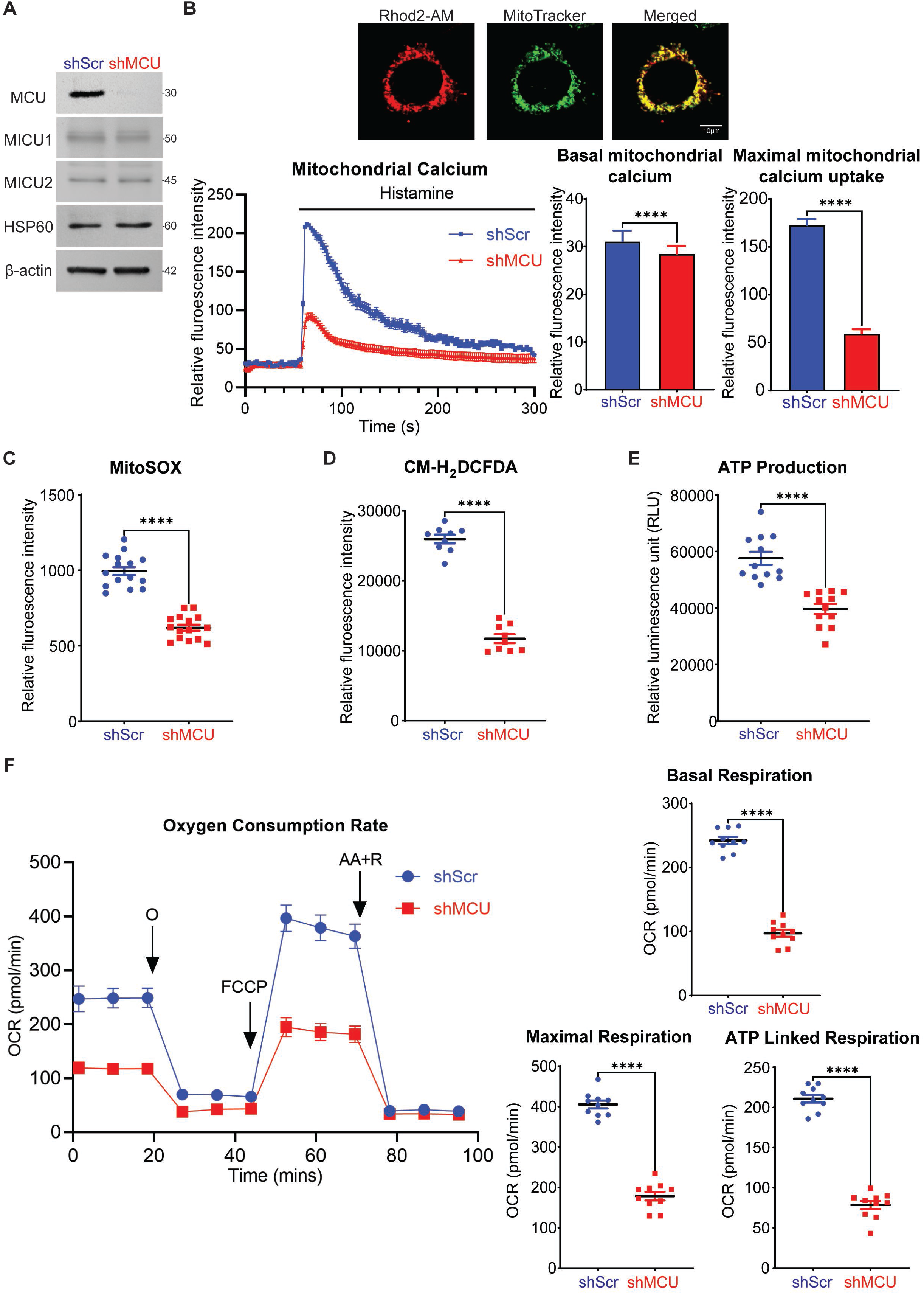
MCU regulates mitochondrial functions in ERMS. A. Western blot analysis showed significant downregulation of MCU expression in stable MCU knockdown RD cells with no change in MICU1, MICU2 and HSP60 expression. The western blot is representative of 3 independent experiments. B. Co-localisation of Rhod2-AM and MitoTracker staining. Scales bar: 10μm. Basal and maximal mitochondrial Ca^2+^ uptake upon induction with 100μM histamine using Rhod2-AM staining is shown in control and shMCU cells (n=3). Values correspond to the average ± SEM. (**** *p*≤0.0001). C. MitoSOX Red staining of shMCU in comparison to shScr is shown (n=5). Values correspond to average ± SEM. (**** *p*≤0.0001). D. Cellular ROS in shMCU as compared to shScr was measured by flow cytometry using CM-H_2_DCFDA staining (n=3). Values correspond to average ± SEM. (**** *p*≤0.0001). E. ATPlite kit revealed reduced ATP production in shMCU as compared to shScr (n=4). Values correspond to average ± SEM. (**** *p*≤0.0001). F. OCR was measured in shMCU cells compared to shScr cells. O: Oligomycin, FCCP: Carbonyl cyanide-4-(trifluoromethoxy)-phenylhydrazone, AA+R: Antimycin A and Rotenone were added accordingly. Values correspond to average ± SEM (n=3). Basal and maximal respiration rate along with ATP production in shScr and shMCU cells is shown. (**** *p*≤0.0001).

### MCU overexpression promotes oncogenic phenotypes

We next examined the phenotype of MCU knockdown cells. A significant reduction in percentage of 5-bromo-2’-deoxy-uridine positive (BrdU^+^) cells was seen in shMCU cells relative to control cells (Fig. 3A). Similarly, transient MCU knockdown in RD, RD18 and JR1 cells reduced their proliferative capacity (Supplementary Fig. 3). RD shScr and shMCU cells were differentiated and stained with anti-myosin heavy chain (MHC) antibody. An increase in MHC^+^ cells was observed in shMCU cells as well as in siMCU RD, RD18 and JR1 cells that was also verified by western blot analysis (Fig. 3B and Supplementary Fig. 4). Myogenin (MYOG), an early myogenic differentiation marker was also elevated (Fig. 3B). We then investigated the migratory and invasive capacity of shScr and shMCU cells. A profound reduction of approximately 80% in the migratory capacity of shMCU and siMCU cells was seen (Fig. 3C and Supplementary Fig. 5). MCU knockdown also significantly decreased invasiveness through matrigel (Fig. 3D).

**Fig. 3.**
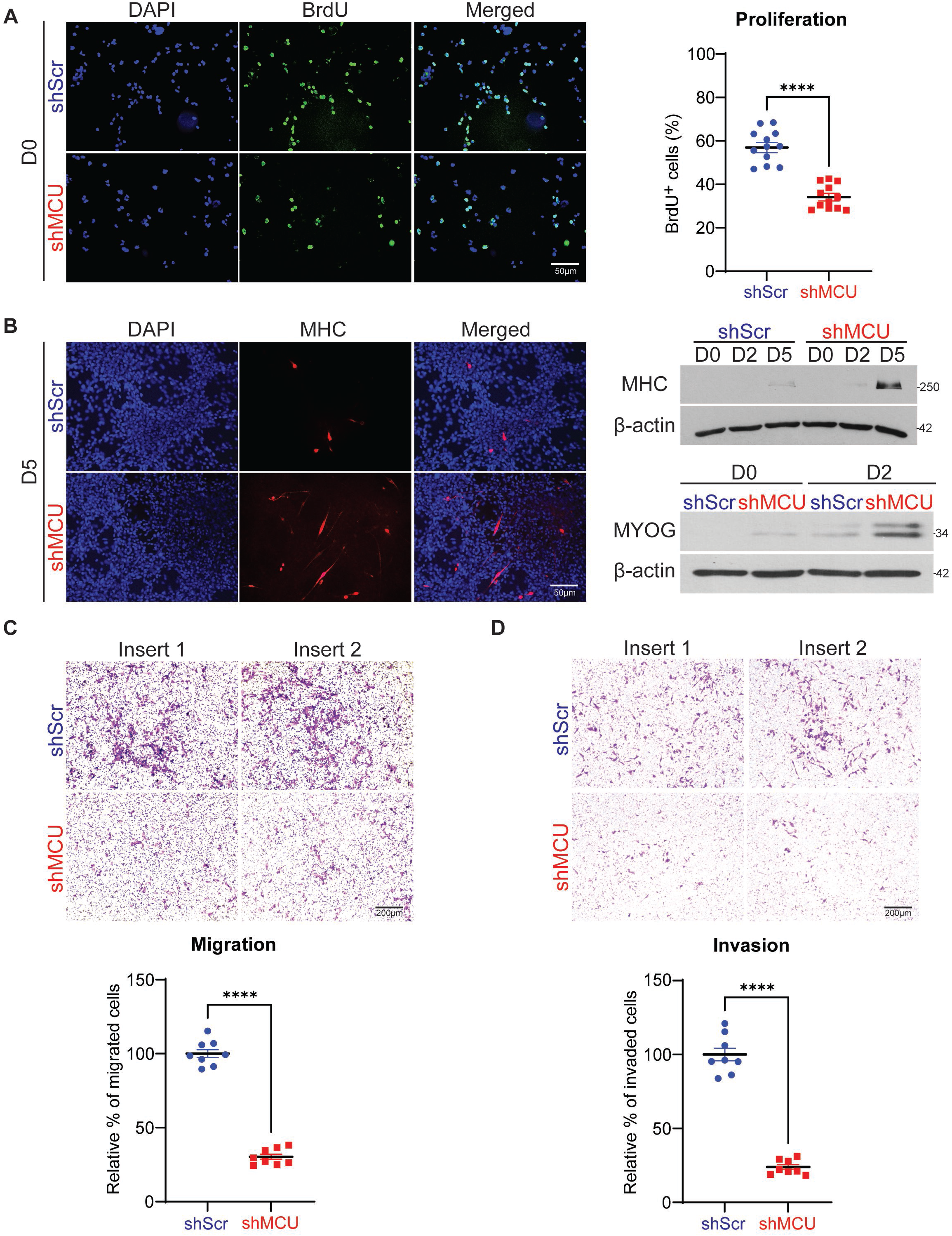
MCU overexpression promotes oncogenic phenotypes. A. BrdU assay to examine proliferation in shScr and shMCU cells. BrdU^+^ cells were analysed by immunofluorescence (n=4). Images are representative of 4 independent experiments. Scale bar: 50μm. The scatter plot shows the percentage of BrdU^+^ cells in shMCU cells relative to controls. The values correspond to average ± SEM. (**** *p*≤0.0001). B. Control shScr and shMCU cells were cultured for 5 days in differentiation medium and analysed by immunofluorescence using anti-MHC antibody. Nuclei were stained with DAPI. Representative images of 4 independent experiments are shown. Scale bar: 50μm. MHC level was analysed in control and shMCU cells by western blot analysis at Day 0 (D0), D2 and D5 in differentiation medium. MYOG level was analysed in control and shMCU cells by western blot analysis at D0 and D2. C. Boyden chamber migration assay of control and shMCU cells. Migrated cells were observed after 24hr using crystal violet staining. Images are representative of 4 independent experiments. Scale bar: 200μm. The relative percentage of migrated cells were quantified in the scatter plot and the values correspond to average ± SEM. (**** *p*≤0.0001). D. Matrigel invasion assay of control and shMCU cells. Invaded cells were stained with crystal violet after 24hr. Images are representative of 4 independent experiments. Scale bar: 200μm. The relative percentage of invaded cells were quantified in the scatter plot and the values correspond to average ± SEM. (**** *p*≤0.0001).

### TGFβ signalling pathway is downregulated upon MCU knockdown

In order to identify mechanisms underlying MCU function, we performed RNA-Sequencing (RNA-Seq). Cluster analysis of differentially expressed genes from control RD and MCU knockdown cells, and volcano plot of differentially expressed genes (Fig. 4A) revealed that 891 genes were significantly up regulated and 1223 genes were significantly down regulated in siMCU cells. Gene Ontology (GO) analysis showed that skeletal system development, muscle organ development and focal adhesion were among the top 5 unique biological processes associated with differentially expressed genes in siMCU cells (Fig. 4B). Kyoto Encyclopaedia of Genes and Genomes (KEGG) pathway analysis identified TGFβ signalling pathway to be among the top 5 significantly altered pathways upon MCU knockdown (Fig. 4C-D). The TGFβ signalling pathway is well known for its role in tumour progression and epithelial to mesenchymal transition (EMT) [31,32]. We therefore focused on examining this pathway. We validated the down regulation of *TGFβ1*, *TGFβR1* and *TGFβR2* expression by qPCR in MCU knockdown cells (Fig. 4E). Moreover, several genes that regulate myogenesis such as myostatin (*MSTN*) [33] and hairy/enhancer-of-split related with YRPW motif protein 2 (*HEY2*) [34] were downregulated, whereas Wnt family member 2B (*WNT2B*) and *WNT5A* were upregulated [35] upon MCU knockdown (Supplementary Fig. 6). Additionally, insulin-like growth factor 2 (*IGF2*) and fibroblast growth factor receptor 4 (*FGFR4*) which are overexpressed in ERMS [2] were down regulated in siMCU cells. Consistent with reduced expression of *TGFβ1*, *TGFβR1* and *TGFβR2*, phosphorylated Smad family member 3 (p-SMAD3) levels, a readout of TGFβ signaling, was reduced in shMCU cells whereas total SMAD3 levels were unchanged (Fig. 4F). Moreover, a significant reduction in the TGFβ reporter 3TP-Lux [36] was seen in shMCU cells (Fig. 4G).

**Fig. 4.**
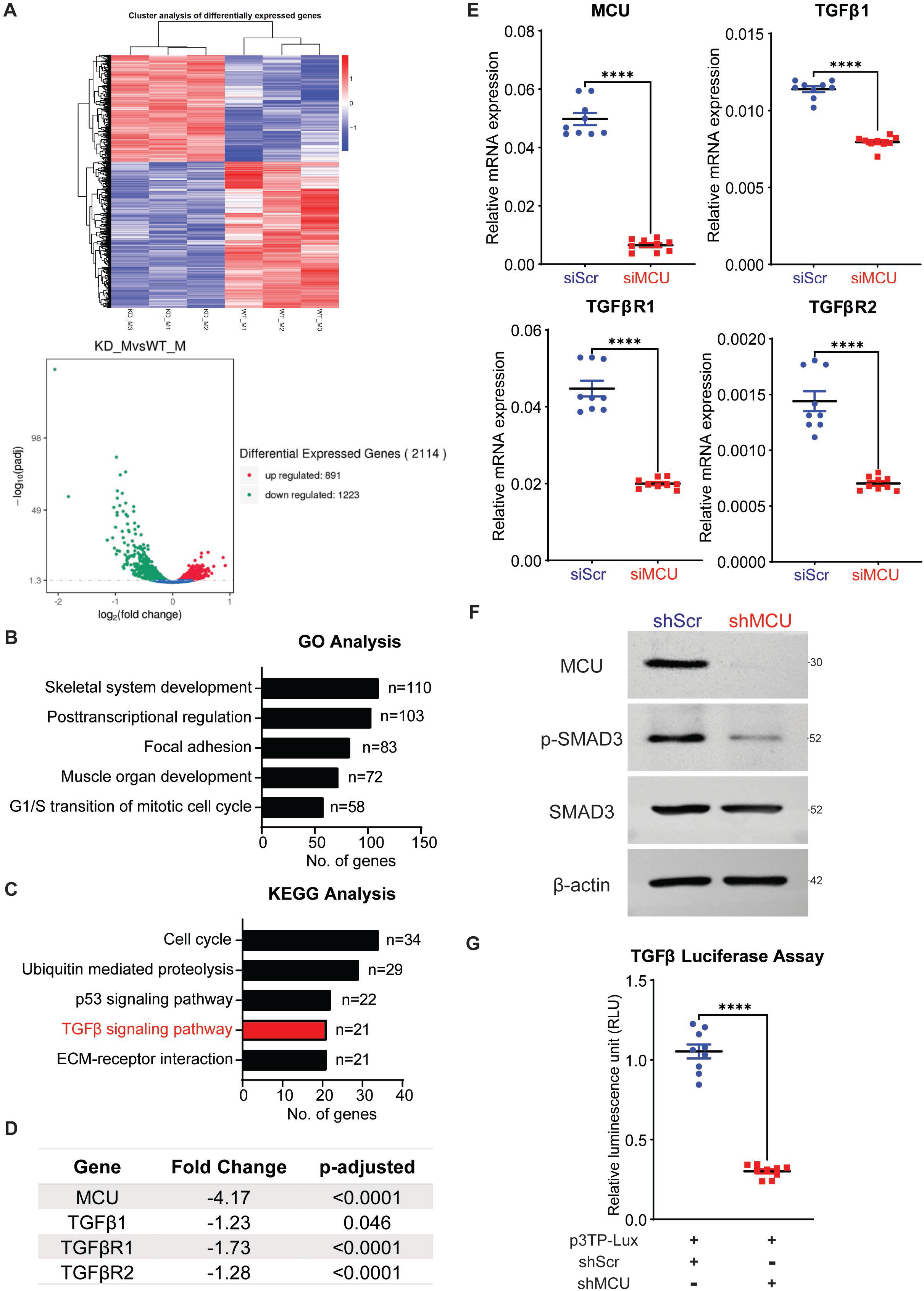
TGFβ signalling pathway is deregulated upon MCU knockdown. A. RNA-Seq heatmap (upper panel) showing hierarchical cluster of differentially expressed genes. Red represents high expression and blue represents low expression. Volcano plot (lower panel) shows distribution of differentially expressed genes upon MCU knockdown. B. GO enrichment histogram showing top five significantly enriched biological processes upon MCU knockdown based on the number of differentially expressed genes. C. KEGG enrichment histogram showing top five unique significantly enriched pathways upon MCU knockdown based on the number of differentially expressed genes. D. A list of the top significantly altered genes in the TGFβ pathway identified by RNA-seq analysis upon MCU knockdown are shown with the fold change and *p* values. E. qPCR analysis for *TGFβ1*, *TGFβR1*, and *TGFβR2* mRNA in control and siMCU cells. The values correspond to average ± SEM (n=3). (**** *p*≤0.0001). F. Western blot analysis of control and shMCU cells using MCU, p-SMAD3 and SMAD3 antibodies. A representative western blot from 3 independent experiments is shown. G. Control and shMCU cells were transfected with 200ng of the TGFβ reporter p3TP-Lux and 5ng Renilla luciferase. Cells were analysed for luciferase activity 48hr later. The values correspond to average ± SEM (n=3). (**** *p*≤0.0001).

### MCU promotes tumour growth in vivo

To examine the impact of MCU loss *in vivo*, we injected RD control (shScr) and shMCU cells in BALB/c nude mice. A significant reduction in tumour growth was apparent in mice injected with shMCU cells (Fig. 5A) without any adverse effect on the mice weight (Fig. 5A, lower panel). Tumour sections from shScr and shMCU cells were analysed histologically and by IHC (Fig. 5B). Ki-67, a proliferation marker was significantly reduced in shMCU tumours. In contrast, myogenic differentiation was strikingly increased as seen from MHC and MYOG levels by IHC and western blot analysis of tumor lysates. Melanoma Cell Adhesion Molecule (MCAM) and Snail Family Transcriptional Repressor 2 (SNAI2), which promote metastasis [37] were decreased. No overt change in active caspase 3 staining was apparent. Moreover, a significant reduction in p-SMAD3 levels was seen in MCU knockdown tumours by western blot analysis (Fig. 5C).

**Fig. 5.**
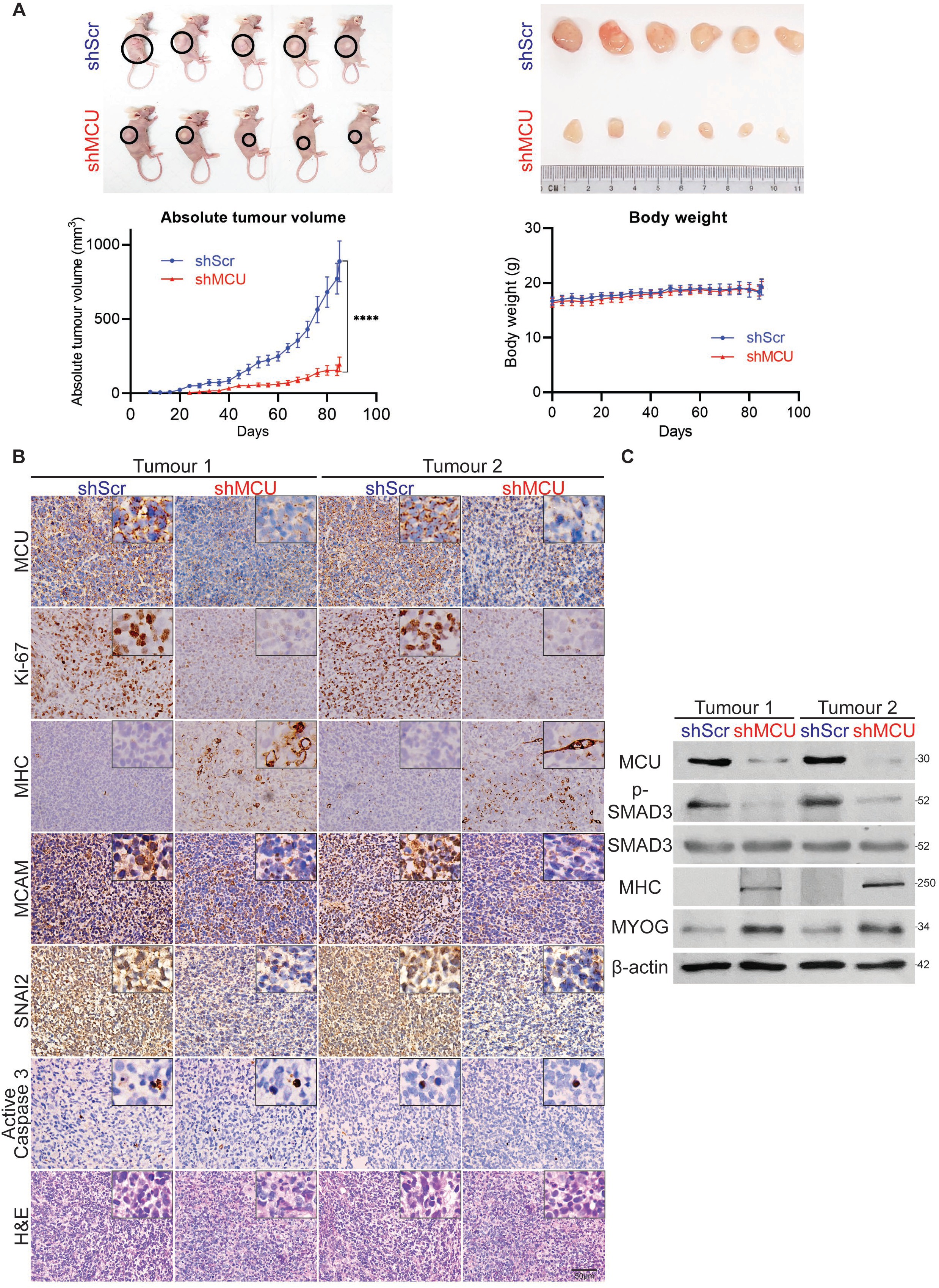
MCU promotes ERMS tumour growth *in vivo*. A. Nude mice were injected with shScr cells (n=7) or shMCU cells (n=7). Representative images of 5 mice in each group (left panel), and resected tumours of 6 mice in each group (right panel) are shown. The absolute tumour volume and body weight are shown in the graphs below. Statistical significance was calculated using repeated-measure one-way ANOVA where **** *p*≤0.0001. Values correspond to the average ± SEM. B. Tumours from two shScr and two shMCU mice were analysed by IHC using anti-MCU, anti-Ki67, anti-MHC, anti-MCAM, anti-SNAI2 and anti-active caspase 3 antibodies. Histology was assessed by haematoxylin and eosin (H&E) staining. Images were taken at 40X magnification. Inset shows 3X zoomed in image. Scale bar: 50μm. C. Two sets of tumour lysates from shScr and shMCU mice were analysed by western blot with anti-p-SMAD3, anti-SMAD3, anti-MHC, anti-MYOG, and anti-β-actin antibodies.

### MCU regulates TGFβ signalling pathway through mROS

Previous studies have shown cross talk between ROS and TGFβ signalling [38,39]. We therefore examined whether mROS was upstream and regulated TGFβ signalling. To alter mROS levels, RD cells were treated with mitoTEMPO (a mROS scavenger) and antimycin A (a complex IV inhibitor) for 48hr. Treatment with mitoTEMPO resulted in a significant reduction in mROS levels in shScr cells, while no futher decrease was observed in shMCU cells. On the other hand, treatment with antimycin A elevated mROS levels in both shScr and shMCU cells. (Fig. 6A). MitoTEMPO treatment of shScr cells significantly reduced TGFβ activity similar to shMCU cells, although shMCU cells showed no futher reduction in TGFβ activity (Fig. 6B). Consistently, p-SMAD3 level was reduced in mitoTEMPO treated shScr cells with no observable difference in mitoTEMPO treated shMCU cells (Fig. 6C). Conversely, treatment of shMCU cells with antimycin A rescued TGFβ reporter activity (Fig. 6B) and p-SMAD3 levels to levels comparable in control DMSO treated cells (Fig. 6C).

**Fig. 6.**
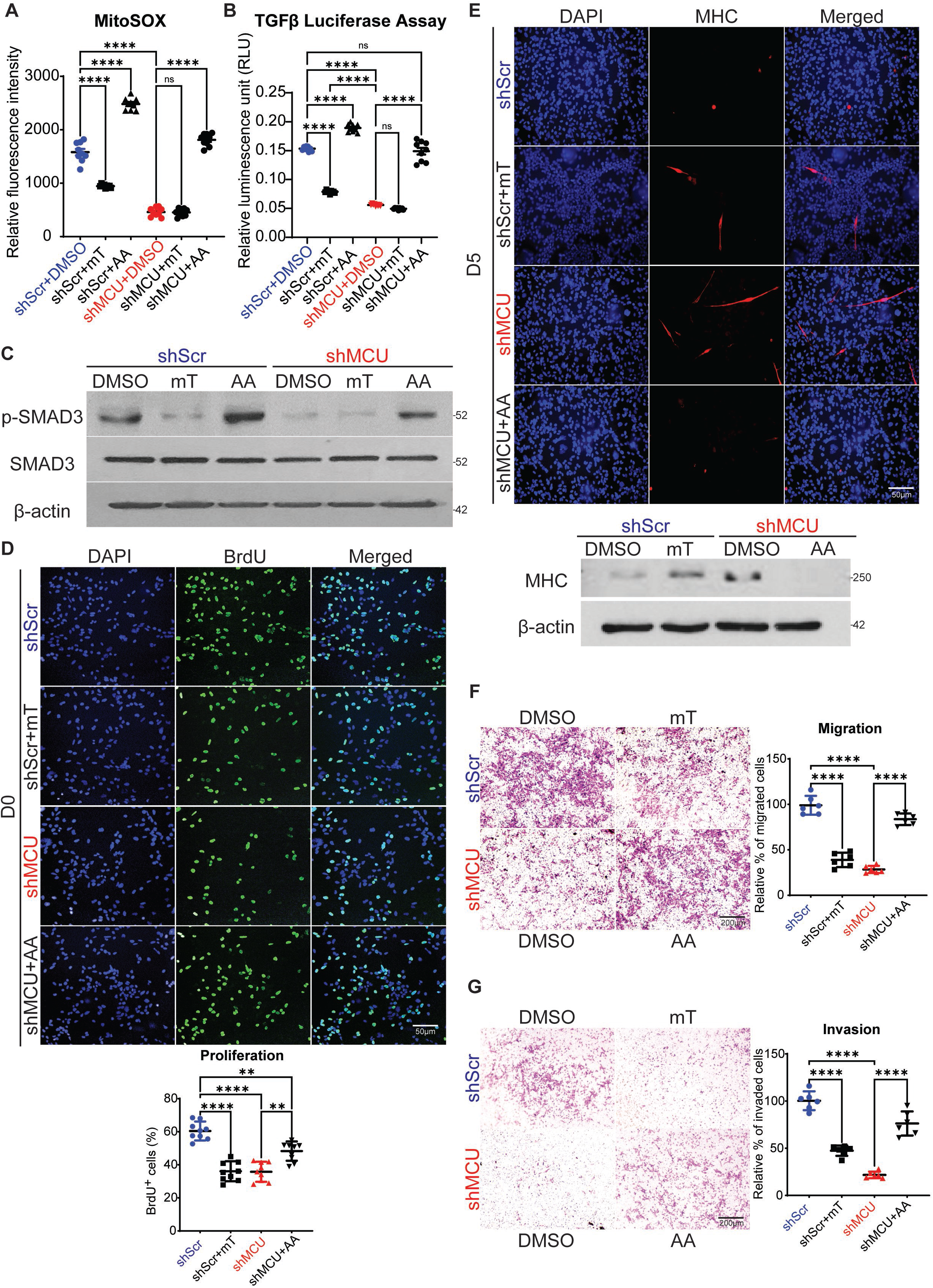
Modulation of mROS impacts TGFβ signalling. A. shScr and shMCU RD cells were treated with DMSO, mitoTEMPO (mT) or antimycin A (AA) for 48hr. MitoSOX staining showed significantly decreased mROS levels upon mT treatment in shScr cells, whereas increased mROS levels were observed with AA treatment in both shScr and shMCU cells. The values correspond to average ± SEM (n=3). (ns=not significant, **** *p*≤0.0001). B. shScr and shMCU cells were treated with mT and AA as indicated for 48hr. Cells were transfected with the p3TP-Lux and analysed for luciferase activity 48hr later. The values correspond to average ± SEM (n=3). (ns=not significant, ** *p*≤0.01, **** *p*≤0.0001). C. p-SMAD3 and SMAD3 levels were examined by western blot analysis in shScr and shMCU cells treated with mT and AA. Representative images of 3 independent experiments are shown. D. Proliferation was analysed by BrdU assay in shScr cells and shMCU cells treated with mT or AA for 48 hr as indicated. Images are representative of 3 independent experiments. Scale bar: 50μm. The bar graph shows the percentage of BrdU^+^ cells in shMCU cells relative to shScr cells. The values correspond to average ± SEM. (** *p*≤0.01, **** *p*≤0.0001). E. shScr cells were treated for 5 days in differentiation media with DMSO or mT and shMCU cells were treated with DMSO or AA. MHC^+^ cells were analyzed by immunofluorescence and quantified using western blot with anti-MHC antibody. Nuclei were stained with DAPI. Representative images of 3 independent experiments are shown. Scale bar: 50μm. F. Migration was analysed for 24hr using Boyden chamber assays following 48hr treatment of shScr cells with DMSO or mT and shMCU cells with DMSO or AA. Images are representative of 3 independent experiments. Scale bar: 200μm. The relative percentage of migrated cells were quantified in the scatter plot. The values correspond to average ± SEM. (**** *p*≤0.0001). G. Matrigel invasion was analysed after 24hr following treatment of shScr cells with DMSO or mT and shMCU cells with DMSO or AA for 48hr. Images are representative of 3 independent experiments. Scale bar: 200μm.The relative percentage of migrated cells were quantified in the scatter plot and the values correspond to average ± SEM. (**** *p*≤0.0001).

To further examined whether the phenotypic effects of MCU depletion are mROS-dependent. Upon treatment of control cells with mitoTEMPO, BrdU^+^ cells were reduced to a level similar to shMCU cells. On the other hand, treatment of shMCU cells with antimycin A partially rescued proliferation (Fig. 6D). Treatment with mitoTEMPO also increased the number of MHC^+^ cells in control cells, and conversely, a prominent reduction in MHC staining was seen upon antimycin A treatment of shMCU cells (Fig. 6E). Similarly, migration and invasion were decreased in mitoTEMPO treated shScr cells, whereas increased migration and invasion were observed in antimycin A treated shMCU cells (Fig. 6F-G). Together, these data demonstrate that modulation of mROS production alters TGFβ signalling and oncogenic phenotypes in ERMS.

## Discussion

The RAS pathway is frequently activated in ERMS and impacts redox balance [6,8,29]. Consistently, ERMS cells are sensitive to drugs that elevate oxidative stress [29]. Despite these correlations, the importance of mitochondrial Ca^2+^ homeostasis has not been examined. Here we show that deregulated expression of the MCU complex impairs mitochondrial Ca^2+^ homeostasis in ERMS cell lines. MCU knockdown caused a reduction in mitochondrial Ca^2+^ uptake and reduced mROS production. This inhibited the TGFβ signalling pathway and impaired proliferation and motility of tumour cells but promoted myogenic differentiation *in vitro* and *in vivo*.

Our finding that MCU positively regulates mitochondrial Ca^2+^ uptake is in concordance with previous studies on MCU knockout mice which show a lack of mitochondrial Ca^2+^ uptake [40,41]. Moreover, an attenuation in mitochondrial Ca^2+^ uptake upon MCU knockdown has been reported in neurons [42], heart [43], liver [15] and pancreatic β cells [44]. The reduced OCR upon MCU knockdown is also in line with similar observations in myofibers of MCU knockout mice [45].

Mitochondria contribute to tumourigenesis and tumour progression in many ways that include the generation of ROS, accumulation of metabolites or alterations in apoptosis [46]. Most of these processes are tightly regulated by Ca^2+^ ions. Since MCU and MICU1 regulate mitochondrial Ca^2+^ uptake and metabolism, deregulation in their expression leads to mitochondrial dysfunction. Indeed, studies have shown increased or decreased MCU and MICU1 expression in different cancers that contributes to tumourigenesis and metastasis in several ways [13,19,20]. In HCC as well as in breast cancer, MCU overexpression results in increased Ca^2+^ uptake and mROS generation which plays an important role in driving tumour progression and metastasis [21,23]. Elevated mROS activates hypoxia-inducible factor 1-alpha (HIF1α), which promotes tumour progression [21]. Additionally, mROS has been reported to reduce superoxide dismutase 2 (SOD2) activity and promote ROS-dependent matrix metalloproteinase (MMP) activity that promotes cell motility [23]. Interestingly, our RNA-Seq data identified a novel signalling pathway downstream of mROS production in ERMS. We show that TGFβ signalling is dampened in response to MCU knockdown. The interplay between ROS and TGFβ signalling pathway has been widely studied [32,38,47,48]. TGFβ signalling is also elevated in ERMS [49,50]. However, increasing or decreasing mROS modulated TGFβ signalling demonstrates that mROS is upstream of TGFβ signalling. The TGFβ pathway has well established roles in cell cycle progression and tumour invasion [31,32]. In addition, TGFβ signalling potently represses myogenic differentiation [49,51]. The impaired TGFβ signalling in shMCU cells correlates with the observed reduction in proliferation and cell motility, with elevated myogenic differentiation *in vitro* and *in vivo*.

In some cancers, MCU overexpression protects cells from apoptosis and thus MCU silencing potentiates cell death [21,52]. While ERMS cells overexpress MCU, we did not observe cell death in shMCU cells *in vitro* and *in vivo*. RNA-Seq analysis showed that the p53 pathway was also altered in response to MCU knockdown. Notably, the expression of several pro-apoptotic genes such as BH3-interacting domain death agonist (*BID*), tumour protein P53 (*TP53*), apoptotic protease activating factor 1 (*APAF1*) and phorbol-12-myristate-13-acetate-induced protein 1 (*PMAIP1*) [53] were reduced significantly upon MCU knockdown. The decreased expression of such pro-apoptotic genes may underlie the absence of apoptosis in shMCU cells.

In addition to MCU overexpression, MICU1 is down regulated in ERMS cell lines. The regulatory mechanisms that underlie these changes in expression are unclear and need further investigation. As shMCU cells showed a modest impact on basal mitochondrial Ca^2+^ levels, it is likely that the down regulation of MICU1 in ERMS cell lines may contribute to the endogenous elevation of basal level of mitochondrial Ca^2+^ [16,17].

Mitochondrial dysfunction is increasingly recognised to have central role in the development of several human diseases including cancer. Pharmacological interventions targeting mitochondria could become effective strategies for treating pathological conditions associated with mitochondrial dysfunction. However, the development of such therapeutic tools is hampered by the incomplete understanding of the molecular mechanisms underlying major mitochondrial functions. In this context, our study elucidates how high MCU expression is linked to tumour progression and a block in myogenic differentiation. Targeting the MCU-mROS-TGFβ axis could be a new unexplored therapeutic strategy in ERMS.

## Materials and methods

### Cell culture, transient and stable knockdown cells

Primary human skeletal muscle myoblasts (HSMM) were purchased from Zen-Bio, Inc. (NC, USA) and cultured in skeletal muscle cell growth medium (#SKM-M, Zen-Bio, USA). Cells were maintained at a confluency of no more than 70% and experiments were performed with cells under 7 passages. RD cells were cultured in Dulbecco’s Modified Eagle Medium (DMEM) (Sigma-Aldrich, St. Louis, MO, USA) with 10% Fetal Bovine Serum (FBS) (HyClone, Cytiva, U.S.) and 1% Penicillin-Streptomycin (HyClone, Cytiva, U.S.). RD18, JR1, RH30 and RH41 were cultured in RPMI 1640 with L-Glutamine (Thermo Fisher Scientific, Waltham, MA, USA) with 10% FBS and 1% Penicillin-Streptomycin. Mouse myoblasts (C2C12) were purchased from ATCC and cultured in DMEM with 20% Fetal Bovine FBSand 1% Penicillin-Streptomycin.

For transient knockdown, cells were transfected with 50nM human MCU-specific siRNA (ON-TARGETplus siRNA SMARTpool, Dharmacon, Lafayette, CO, USA) containing a pool of three to five 19-25 nucleotide siRNAs. Control cells were transfected with 50nM scrambled siRNA (ON-TARGETplus, non-targeting pool, Dharmacon, Lafayette, CO, USA). Transfections were performed using Lipofectamine RNAiMax (Thermo Fisher scientific). Cells were analysed 48hr post-transfection for all assays.

For generation of stable knockdown cell lines, HEK293FT cells were transfected with packaging plasmid pIP1 (5μg) and pIP2 (5μg), envelope plasmid pIP/VSV-G (5μg) (ViraPower™ Lentiviral Packaging Mix, Thermo Fisher Scientific) and 5μg lentiviral expression constructs shRNA (pLKO.1 Mission shRNA DNA clone, Sigma-Aldrich Inc.) or shMCU (#SHCLNG-NM_138357 Mission shRNA, Sigma-Aldrich Inc.). 16hr post-transfection, the cell supernatant was replaced with basal DMEM medium. The supernatants were centrifuged, and the viral pellet was resuspended in DMEM medium. RD cells were transduced with shRNA control lentivirus particles or shMCU lentivirus particles with polybrene (8μg/ml) (Sigma-Aldrich Inc.). Transduced cells were selected with 1μg/ml puromycin (Sigma-Aldrich Inc.) for three days until all cells in control plates were dead.

### Mitochondrial calcium measurement

Cells were plated on glass bottom dishes and loaded with 5μM Rhod-2 AM (Sigma-Aldrich Inc.) and 100nM MitoTracker Green FM (Invitrogen) in extracellular medium as described previously [54,55]. Cells were incubated for 50min at 37°C before washing with the same extracellular buffer containing 0.25% BSA at room temperature for 20min. To measure mitochondrial Ca^2+^, the dishes were mounted on an on-stage incubator at 37°C with 5% CO_2_ and imaged with confocal microscope with 60X water objective lens. After 1min of baseline recording, 100μM histamine (Sigma-Aldrich Inc.) was added to induce mitochondrial Ca^2+^ uptake. Confocal images were recorded every 1s at 561nm excitation for another 4min. The fluorescence intensities of the images were analysed and quantified with Image J (NIH). Mitochondrial Ca^2+^ changes were quantified by plotting relative fluorescence intensity of the images for a duration of 5min. Basal mitochondrial Ca^2+^ was quantified by measuring relative fluorescence intensity during the first 1min of baseline recording. Maximal mitochondrial Ca^2+^ uptake was quantified by the difference between maximal fluorescence intensity and basal fluorescence intensity.

### Reactive oxygen species

Cellular reactive oxygen species (ROS) and mitochondrial ROS (mROS) were detected using fluorescence probe CM-H_2_DCFDA (Invitrogen; Thermo Fisher Scientific, Inc., USA) and MitoSOX Red (Invitrogen; Thermo Fisher Scientific, Inc., USA) respectively. Cells were trypsinised and loaded with 5μM CM-H_2_DCFDA or 5μM MitoSOX Red for 20min at 37°C respectively. Fluorescence intensity was analysed using flow cytometry. A minimum of 100,000 events per sample were collected and the data was analysed using CytExpert software (Beckman Coulter, Inc.). To modulate mROS levels, RD cells were treated with 200nM of mitoTEMPO (mT), whereas shMCU cells were treated 500nM antimycin A (AA) for 48hr. DMSO was used as a control.

### ATP measurement

ATP production was measured with the ATPlite Luminescence Assay System (PerkinElmer) according to manufacturer’s instructions.

### Oxygen consumption rate measurement

Oxygen consumption rate (OCR) was measured with a XF24 extracellular analyser (Seahorse Bioscience) and XF Cell Mito Stress Test Kit (Seahorse Bioscience). Cells were seeded at 50,000 cells/well (~80-90% confluent when assayed) in a 24-well Agilent Seahorse XF Cell Culture Microplate (Seahorse Bioscience) and incubated overnight at 37°C. Prior to the assay, growth media was replaced with XF DMEM medium, pH7.4 (Seahorse Bioscience) supplemented with 1mM sodium pyruvate (Sigma-Aldrich, St. Louis, MO, USA) and 10mM glucose (Sigma-Aldrich, St. Louis, MO, USA). Cells were then incubated for 45min to 1hr in 37°C without CO_2_ to prevent acidification of medium. After loading the plate into the machine, basal respiration rate was measured before cells were exposed sequentially to oligomycin (1μM), carbonyl cyanide p-trifluoromethoxyphenylhydrazone (FCCP; 1μM) and rotenone + antimycin A (500nM each). After each injection, OCR was measured for 5min, the medium was mixed and again measured for another 5min. After the experiment, protein concentration was determined by lysing samples in each well and performing Bradford analysis (Bio-Rad). Maximum respiration rate was quantified by maximal OCR after adding FCCP. ATP production was quantified by the decrease in OCR upon injection of the ATP synthase inhibitor oligomycin.

### Reporter assays

TGFβ reporter assay was analysed as described [56]. Briefly, shScr and shMCU cells were transfected with 200ng of 3TP-Lux reporter in 24-well plates. 5ng of Renilla reporter was co-transfected as an internal normalisation control. Transfection was carried out in triplicates using Lipofectamine 3000 Transfection Reagent (ThermoFisher Scientific). Reporter activity was analysed with the Dual-Luciferase Reporter Assay System (Promega). Luminescence was analysed with Varioskan plate reader using the SkanIT software.

### Western blot analysis

Whole cell and tumour lysates were isolated using RIPA buffer supplemented with protease inhibitors (Complete Mini, Sigma-Aldrich Inc.) and phosphatase inhibitors including sodium pyrophosphate, β-glycerophosphate, sodium fluoride and sodium orthovanadate (Sigma-Aldrich). The following primary antibodies were used: anti-MCU (#D2Z3B 1:1000, Cell Signalling), anti-MICU1 (#HPA037479 1:1000, Sigma-Aldrich), anti-MICU2 (#ab101465 1:1000, Abcam), anti-phospho-SMAD3 (#C25A9, 1:1000, Cell Signalling), anti-SMAD3 (#9513, 1:1000, Cell Signalling), anti-MYOG (#sc-12732, 1:500, Santa-Cruz), anti-MHC (#sc-32732, 1:250, Santa-Cruz) anti-HSP60 (#611563, BD Biosciences) and anti-β-actin (#A2228, 1:10,000, Sigma-Aldrich). Appropriate secondary antibodies (IgG-Fc Specific-Peroxidase) of mouse or rabbit origin (Sigma-Aldrich) were used.

### RNA sequencing (RNA-Seq) and quantitative real-time polymerase chain reaction (q-PCR)

For RNA-Seq analysis, RNA was isolated from control (siScr) and siMCU cells in triplicates using Trizol. RNA purity and integrity were assessed with Nanodrop, agarose gel electrophoresis and Agilent 2100. Raw image data file from Illumina (HiSeq PE150) was transformed to Sequenced Reads by CASAVA base recognition and stored in FASTQ(fq) format. Raw reads were filtered in order to achieve clean reads using the following filtering conditions: reads without adaptors, reads containing number of base that cannot be determined below 10% and at least 50% bases of the reads having Qscore denoting Quality value <=5. For mapping of the reads, STAR software was used. 1M base was used as the sliding window for distribution of the mapped reads. For analysis of differentially expressed genes, Gene Ontology (GO) and Kyoto Encyclopaedia of Genes and Genomes (KEGG) analysis were done with corrected *p*-value less than 0.05 as significant enrichment.

For q-PCR analysis, total RNA was extracted using Trizol (Thermo Fisher Scientific) and quantified using Nanodrop. Messenger RNA (mRNA) was converted to a single-stranded complementary DNA (cDNA) using iScript cDNA Synthesis Kit (Bio-Rad). Q-PCR was performed using Lightcycler 480 SYBR Green 1 Master Kit (Roche). PCR amplification was performed as follows: 95°C 5min, followed by 95°C for 10s, annealing at 60°C for 10s, and 45 cycles at 72°C for 10s. Light Cycler 480 software (version 1.3.0.0705) was used for analysis. CT values of samples were normalized to internal control GAPDH to obtain delta CT (ΔCT). Relative expression was calculated by 2^−ΔCT^ equation. q-PCR was done using technical triplicates and at least three independent biological replicates were done for each analysis. Representative data is shown. Primer sequences can be found in Supplementary Table 1. RNA-seq data has been deposited in GEO under the accession number GSE173200 and can be viewed with the token: gbexwwkifjgrdip

### Proliferation and differentiation assays

Proliferation and differentiation were analysed as described [57, 58]. Briefly, proliferation was measured using 5-bromo-2’-deoxy-uridine (BrdU) labelling (Roche, Basel, Switzerland). Cells were pulsed with 10μM BrdU, fixed and incubated with anti-BrdU antibody (1:100) followed by anti-mouse Ig-fluorescein antibody (1:200) and mounted onto a glass slide using DAPI (Vectashield, Vector Laboratories, CA, USA). Images were captured using fluorescence microscope BX53 (Olympus Corporation, Shinjuku, Tokyo, Japan) at 40X magnification.

For differentiation, cells were cultured in differentiation media consisting of either basal DMEM or RPMI 1640 with 2% Horse Serum (HyClone, Cytiva, U.S.) for 2-5 days. Cells were incubated with anti-MHC primary antibody (MHC; R&D Systems, Minneapolis, MN, USA) (1:400) followed by secondary goat anti-Mouse IgG (H+L) Highly Cross-Adsorbed Secondary Antibody, Alexa Fluor 568 (Thermo Fisher scientific). Coverslips were mounted with DAPI (Vectashield, Vector Laboratories, CA, USA) and imaged with BX53 microscope (Olympus Corporation) at 40X magnification.

### Migration and invasion assay

Migratory and invasive capacity were assessed as described [57,58] with Boyden chamber (Greiner Bio-One). Briefly, cells were serum deprived for at least 12hr and seeded at a density of 50,000 cell per well in serum-free media. 10% FBS containing media was added to the lower chamber. The inserts were stained with crystal violet after 24hr and imaged at 10X magnification. The invasive capacity of the cells was determined similarly using inserts coated with matrigel (Bio Lab) and cells were seeded at a density of 70,000 cells per insert.

### Mouse xenograft experiments

6-week-old C.Cg/AnNTac-Foxn1^nu^NE9 female BALB/c nude mice (InVivos, Singapore) were injected subcutaneously in the right flank with either shScr or shMCU RD cells (10^6^ cells per mice). 7 mice were used in each group. The number of mice per group was determined using power analysis assuming 5% significance level and 80% statistical power with 10% attrition rate. Tumour onset and growth were monitored every alternate day. Tumour diameter was measured, and volume was calculated using the following formula: V=(LxWxW)/2, where V=tumour volume, L=tumour length, W=tumour width. Resected tumours were used to prepare tumour lysates for western blot analysis or fixed with formalin for histopathological analysis. All animal procedures were approved by the Institutional Animal Care and Use Committee under the protocol number R19-0890.

### Immunohistochemistry (IHC)

Paraffin sections of 6 archival primary ERMS tumours from KK Women’s and Children Hospital in Singapore were analysed by IHC using anti-MCU antibody (1:50, Sigma-Aldrich). Following Institutional Review Board approval (CIRB 2014/20179), specimens were obtained from patients at KK Women’s and Children Hospital who were recruited prospectively, with written parental consent and child assent obtained. TMA (SO2082b), comprising of 27 ERMS tumour specimens and 8 striated muscle tissue, was purchased from US Biomax, Inc. and analysed by IHC using anti-MCU antibody (1:50, Sigma-Aldrich) following the manufacturer’s protocol. Paraffin sections from mouse xenografts were stained with haematoxylin and eosin and analysed by IHC as described [57,58]. Sections were incubated overnight at 4°C with anti-MCU (1:50, Sigma-Aldrich), anti-Ki67 (1:100, Santa Cruz Biotechnology), anti-MCAM (1:200, Proteintech), anti-SNAI2 (1:100, Proteintech), anti-active caspase 3 (1:200, Cell Signalling), anti-MHC (1:200, Santa Cruz Biotechnology) antibodies using Dako REAL EnVIsion-HPR, Rabbit-Mouse kit (Dako, Denmark). Sections were counterstained with haematoxylin (Sigma-Aldrich). Slides were dehydrated and mounted using DPX (Sigma-Aldrich) and imaged using BX53 Olympus microscope at 40X magnification.

### Statistical analysis

For statistical analysis, two-tailed non-parametric unpaired t-test was used to evaluate the significance between data sets with the use of GraphPad prism 9.0 software. For animal xenograft experiment and rescue experiments, one-way ANOVA test with appropriate correction was performed with GraphPad prism 9.0 software. Each experiment was performed thrice as independent biological replicates. Each independent experiment had three technical replicates with the exception of migration and invasion assay which had two technical replicates each. All technical replicates were plotted on the scatter plots. Standard error of mean was calculated for all data sets and a *p*-value less than 0.05 was considered statistically significant.

## Acknowledgement

We thank Peter Houghton (Nationwide Children’s Hospital, Ohio, USA) and Rosella Rota (Bambino Gesu Children’s Hospital, Rome, Italy) for the ERMS cell lines, Jeff Wrana (Lunenfeld-Tanenbaum Research Institute, Mount Sinai Hospital, Toronto, Canada) for 3TP-Lux reporter and Karthik Mallilankaraman (National University of Singapore) for valuable advice, discussions and imaging buffers. This work was supported by the Ministry of Education grant [NUHSRO/2020/149/T1/Seed-Sep/03] to RT, VIVA Foundation for Children with Cancer (VIVA-KKH Paediatric Brain and Solid Tumour Programme) to AHPL. HYC is supported by the President’s Graduate Scholarship at the National University of Singapore.

## Author contributions

HYC did all the experiments. AL provided tumor samples and analyzed the patient IHC data. HYC and RT analysed the data and wrote the manuscript.

## Data Availability

The RNA-seq data has been deposited in GEO under the accession number GSE173200 and can be viewed with the token: gbexwwkifjgrdip

